# A general *in vitro* assay to study enzymatic activities of the ubiquitin system

**DOI:** 10.1101/660894

**Authors:** Yukun Zuo, Boon Keat Chong, Kun Jiang, Daniel Finley, David Klenerman, Yu Ye

## Abstract

The ubiquitin (Ub) system regulates a wide range of cellular signaling pathways. Several hundred E1, E2 and E3 enzymes are together responsible for protein ubiquitination, thereby controlling cellular activities. Due to the numerous enzymes and processes involved, studies on ubiquitination activities have been challenging. We here report a novel FRET-based assay to study the *in vitro* kinetics of ubiquitination. FRET is established between binding of fluorophore-labeled Ub to eGFP-tagged ZnUBP, a domain that exclusively binds unconjugated Ub. We name this assay the Free Ub Sensor System (FUSS). Using Uba1, UbcH5 and CHIP as model E1, E2 and E3 enzymes, respectively, we demonstrate that ubiquitination results in decreasing FRET efficiency, from which reaction rates can be determined. Further treatment with USP21, a deubiquitinase, leads to increased FRET efficiency, confirming the reversibility of the assay. We subsequently use this assay to show that increasing the concentration of CHIP or UbcH5 but not Uba1 enhances ubiquitination rates, and develop a novel machine learning approach to model ubiquitination. The overall ubiquitination activity is also increased upon incubation with tau, a substrate of CHIP. Our data together demonstrate the versatile applications of a novel ubiquitination assay that does not require labeling of E1, E2, E3 or substrates, and is thus likely compatible with any E1-E2-E3 combinations.

## Introduction

Ubiquitination is a posttranslational modification process that dictates the destiny of a wide range of proteins, thereby regulating many aspects of cellular functions^1^. The typical ubiquitination reaction involves the sequential activity of three enzymes, E1, E2 and E3, which together conjugate the C-terminal carboxyl group of ubiquitin (Ub) to the ε-amino group of a Lys residue of a target protein^2^. The C-terminus of Ub is first activated by the E1 in an ATP-dependent manner and charges the Ub to its active site Cys, resulting in a thioester bond between the enzyme and Ub (E1∼Ub)^3^. There are ∼40 known E2s, which can accept Ub to their active site Cys through a thioester transfer reaction from the E1^4,5^. Charged E2∼Ub can subsequently transfer the Ub to the active site Cys of HECT family E3s, which will identify and ubiquitinate substrates directly^6^. Alternatively for RING family E3s, reactions are catalyzed by bringing together the substrate and the charged E2 in an optimal orientation for ubiquitination^7^. Of the >700 E3s, the majority are RING E3s or related to RING, such as Cullin E3s, which provide a multisubunit scaffolding mechanism for catalysis^8^; RBR (RING-Between-RING) E3s^9^, which can receive charged Ub much like a HECT E3; and U-box domain E3s^10^, which are structurally similar but achieve catalysis without the two Zn^2+^ ions critical for the activity of canonical RING domains. A number of deubiquitinases (DUBs) also exist in cells to remove Ub modifications, thereby acting as negative regulators of ubiquitination^11^.

Ubiquitinated substrates can be modified on either a single or multiple Lys residues, known as mono- or multiubiquitination, respectively^12^. The Ub itself may also become ubiquitinated, forming distinct Ub chains (polyubiquitination) on the substrate^13^. The complexity of ubiquitination is further increased by branched or forked Ub chains, where a Ub moiety already part of a chain is further ubiquitinated on a second residue^14^. Moreover, Thr and Ser have also been reported to be ubiquitinated as part of the ERAD^52^ or viral infection mechanism^15^. A recent study has identified and characterized the mechanisms by which MYCBP2, a RING E3, preferentially ubiquitinates Thr over Ser residues on substrates^16^. The Ub system is therefore regulated through layers of complex enzymatic reactions resulting in a wide range of possible ubiquitination types.

The distinction with which ubiquitination activities regulate cellular functions makes enzymes of this system interesting targets for therapeutic intervention^17,18^. A number of experimental assays have been developed to study DUBs (e.g. ^19–22^). In comparison, few assays are available to study the concerted action of E1, E2 and E3s. Quantification of ubiquitination rates has previously relied on Western blots (e.g. ^23–25^), a time-consuming and expensive process. Assays based on fluorescence polarization have also been developed to exploit changes in the mass, and hence the tumbling rate, of dye-labeled Ub in free versus conjugated state^25,26^. Other commercially available kits, such as Lanthascreen^26^, depend on the availability and specificity of the antibody against the substrate; or in the case of AMP-Glo^27^, measures ubiquitination activity indirectly through the rate of ATP hydrolysis, where the activities of E2 and E3 are not directly involved. A recent work has exploited mass spectrometry to study ubiquitination^28^. While sensitive, this method is likely to be expensive as it requires both specialist staff time and access to a MALDI-TOF instrument. There is currently a need for a simple and inexpensive assay to study real-time ubiquitination and deubiquitination events directly using canonical instruments typically found in biochemistry laboratories.

In this study, we describe a novel Förster resonance energy transfer (FRET) assay that enables sensitive measurement of ubiquitination in real time. FRET was established between fluorophore-labeled Ub and an eGFP-tagged ZnUBP domain that specifically binds free but not conjugated Ub. Ubiquitination catalyzed by the Uba1-UbcH5-CHIP cascade reduced the FRET efficiency, while subsequent deubiquitination by USP21 restored the FRET. Enzyme kinetics could be determined from initial rates of reaction measured with FRET. We systematically varied the concentration of Uba1, UbcH5 or CHIP over a wide range and found that increasing the concentration of UbcH5 or CHIP but not Uba1 enhanced the rate of ubiquitination. Together, we describe a novel assay that directly reports enzymatic activity of the ubiquitin system and is optimized for measurements in 96-well plates in a high-throughput manner. As this assay does not require modification or labeling of enzymes involved, it is likely applicable to a wide range of Ub-interacting partners (UbIPs) that catalyze ubiquitination and deubiquitination.

## Materials and Methods

### Molecular Biology

ZnUBP domain (encoding residues 174-289 of human USP5/IsoT) was cloned into a pOPINGFP vector resulting in the DNA sequence of eGFP fused in frame to the 3’-end of ZnUBP (kindly shared by David Komander). The ubiquitin construct in pOPINS vector was modified at 5’-end to include codons for Cys and Ala immediately before the start of the first Met^22^. The resulting construct translated into a fusion protein His_6_-SUMO-Cys-Ala-ubiquitin. Plasmids for full-length mouse CHIP in pGEX-6P vector (kind gift from Sophie Jackson), full-length mouse Uba1 in pET28a, human UbcH5a in pGEX-6P and human USP21^29^ in pOPINS vector were used for protein expression (kind gifts from David Komander).

### Protein purification

Plasmids were transformed into BL21 cells and grown at 37°C shaking conditions in LB media until O.D. >1.0. At this cell density cultures were induced with 1 mM IPTG and incubated overnight at 20°C. Subsequently, cells were collected by centrifugation at 4800 r.p.m. using a JLA-8.1000 rotor on an Avanti J-26 XP (Beckman, Palo Alto, USA). Cell pellets were resuspended in 40 mL of Lysis buffer (200 mM NaCl, 10 mM DTT, 25 mM Tris, pH 8.0) for GST-tagged proteins, or Buffer A (300 mM NaCl, 10 mM imidazole, 50 mM Tris, pH 7.4) for His_6_-tagged proteins. Cell lysis was achieved by sonication and the lysate was cleared by centrifugation at 20,000 rpm for 30 min using a pre-chilled JA-25.50 rotor at 4°C on Avanti J-26 XP.

For affinity purification of GST-tagged proteins, the cleared cell lysate was filtered through a 0.45 μm filter and incubated with 2-3 mL glutathione beads in a glass column for 1 hr under constant agitation at 4°C. Subsequently, the beads were washed with 1 L of High salt buffer (500 mM NaCl, 5 mM DTT, 25 mM Tris, pH 8.5) and 1 L of Low salt buffer (50 mM NaCl, 5 mM DTT, 25 mM Tris, pH 8.5) to remove non-specifically bound proteins. PreScission protease was then added to the beads in Low salt buffer and incubated overnight at 4°C. On the next day, the protein of interest was eluted from the beads. Eluted fractions that were more than 90% pure on Coomassie-stained protein gels were pooled and concentrated to >1 mg/mL and flash-frozen.

For affinity purification of His_6_-tagged proteins, the cleared cell lysate was filtered through a 0.22 μm filter and loaded onto a pre-packed TALON column pre-charged with cobalt ions. Unbound proteins were washed out with 50 mL of Buffer A and the protein of interest was eluted using Buffer A + 200 mM imidazole (pH adjusted to 7.4). The eluted protein fractions were pooled and dialyzed against Low salt buffer overnight at 4°C. Proteins expressed from pOPINS vectors were incubated with SENP1 protease during the dialysis to remove the N-terminal His_6_-SUMO tag.

USP21 and Ub^-1Cys^ were further purified by pre-packed Resource-S and Mono-S (GE Healthcare) cation-exchange chromatography columns, respectively, with a linear salt gradient reaching 0.5 M NaCl. ZnUBP^eGFP^ was loaded onto Resource-Q anion-exchange column running the same linear gradient. All three proteins eluted very early from the column and fractions containing the protein of interest were concentrated to 2-5 mL and loaded onto a Superdex75 gel-filtration column in Protein buffer (50 mM NaCl, 50 mM Tris, 1 mM DTT, pH 7.4). Fractions from the major peak containing the protein of interest were pooled together, concentrated and assessed on SDS-PAGE to be pure before being flash-frozen at >1 mg/mL concentration. All column chromatography methods were performed at 4°C on an AKTA Purifier system.

Full-length tau (isoform 0N4R) was purified as described before^30^.

### Protein labeling

AlexaFluor-594 C5 maleimide and AlexaFluor-647 C2 maleimide were dissolved in DMSO (10 mM final concentration), flash-frozen in 20 µL aliquots and stored at -80°C. Labeling of Ub^-1Cys^ was achieved by reacting ∼300 μM of Ub in Labeling buffer (1 mM TCEP, 50 mM Tris, pH 7.2) with 1.2-fold molar excess of fluorophore. The reaction mixture was agitated in the dark at room temperature for three hours. Unreacted fluorophore was removed by size-exclusion chromatography (HiLoad S26/10 column). Labeled proteins were concentrated to > 200 µM concentration as measured on nanodrop.

### FUSS and ubiquitination reactions

ZnUBP^eGFP^ and Ub^594^ were diluted in Protein buffer down to the desired concentration. The final concentration of ZnUBP^eGFP^ was 0.1 µM in all the reactions. For cuvette-based titration experiments, Ub^594^ was serially diluted in Protein buffer containing ZnUBP^eGFP^ at 0.1 µM so as to keep the final ZnUBP^eGFP^ concentration constant.

All reactions were performed in Protein buffer. ZnUBP^eGFP^ and Ub^594^ were mixed with Uba1, UbcH5 and CHIP and incubated for 5 min to allow equilibration prior to reaction initiation. The reaction volume for cuvette-based experiments was 50 µL and 20 µL per well for 96-well plates. Ubiquitination reactions were initiated by adding stock 250 mM ATP-MgCl_2_ (pH-adjusted to 7.0) to a final concentration of 10 mM in the reaction.

### Fluorescence measurements

Cary Eclipse spectrophotometer (Varian Inc.) was used for cuvette-based fluorescence measurements. The fluorescent sample was loaded in a quartz cuvette (Hellma) with 3.0 × 10.0 mm windows placed so that detection would be perpendicular to the excitation light path. For fluorescence emission scans the excitation maximum wavelength was set to 488 nm with ± 5 nm window. PMT was set (700 V) so that the emission maximum lies approximately half-way between the minimum and maximum fluorescence detection limits. Donor and acceptor intensities were calculated from the integral of the spectrum between 500 to 560 nm and 610 to 670 nm, respectively. The interval time between measurements was set to 1 min unless otherwise indicated.

Clariostar plate-reader (BMG Labtech) was used for simultaneous fluorescence measurements of multiple reactions. Each reaction contained 20 µL of sample volume in black 96-well round-bottom plates (Corning Life Sciences). The focal height was adjusted to the fluorescence intensity of the most concentrated sample. Each well was excited with 10 flash pulses and measured for 200 cycles with 60 s per cycle. The fluorescence gain for the donor (ex_440-10_) and the acceptor (em_610-10_) was set to 2000, unless otherwise indicated. The wells were covered with a sealable aluminum foil and a small amount of water was added between the wells to reduce evaporation during data recording.

### Fluorescence scans of protein gels and Western blots

Reaction samples were taken and quenched with LDS buffer containing reducing agent (Invitrogen) in a 2:1 volumetric ratio (sample:LDS) unless otherwise stated. Proteins were separated on NuPAGE Bolt 4-12% SDS-polyacrylamide gels (Invitrogen) and subsequently scanned on a Typhoon scanner using the indicated excitation and emission modules. A Trans-Blot (Biorad) system was used to transfer proteins separated by SDS-PAGE to PVDF membranes. Primary rabbit monoclonal anti-CHIP (EPR4447, Abcam) or mouse monoclonal anti-tau (1E1/A6, Millipore) were recognized by secondary anti-rabbit or anti-mouse antibodies carrying AlexaFluor-647 labels (Invitrogen) and detected on the Typhoon scanner using the same module as above.

### Data analysis

All data analyses were carried out on Prism 6 (GraphPad) unless otherwise indicated. FRET efficiency values reported in this manuscript are calculated using the equation:

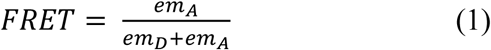

where *em*_*D*_ and *em*_*A*_ are emission intensities of the donor and acceptor, respectively.

The binding constant, *K*_*D*_, describes the relationship between bound and unbound ZnUBP^eGFP^ and is defined by the equation below:

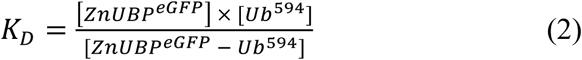

where [ZnUBP^eGFP^] and [Ub^594^] are the concentrations of the free protein and ligand, respectively, and [ZnUBP^eGFP^ – Ub^594^] that of the bound complex.

Experimentally, the binding constant is derived from data extrapolation by applying the following equation:

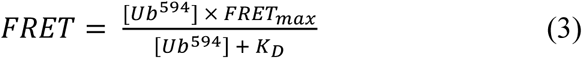

Cuvette-based FRET measurements are converted to Ub^594^ concentrations using the function from **Figure 1d**:

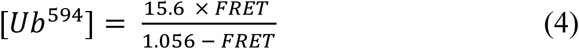

and that derived from **Figure S1** to convert FRET measured on the plate-reader:

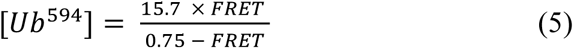

where the Ub^594^ concentration is in µM.

**Figure 1.**
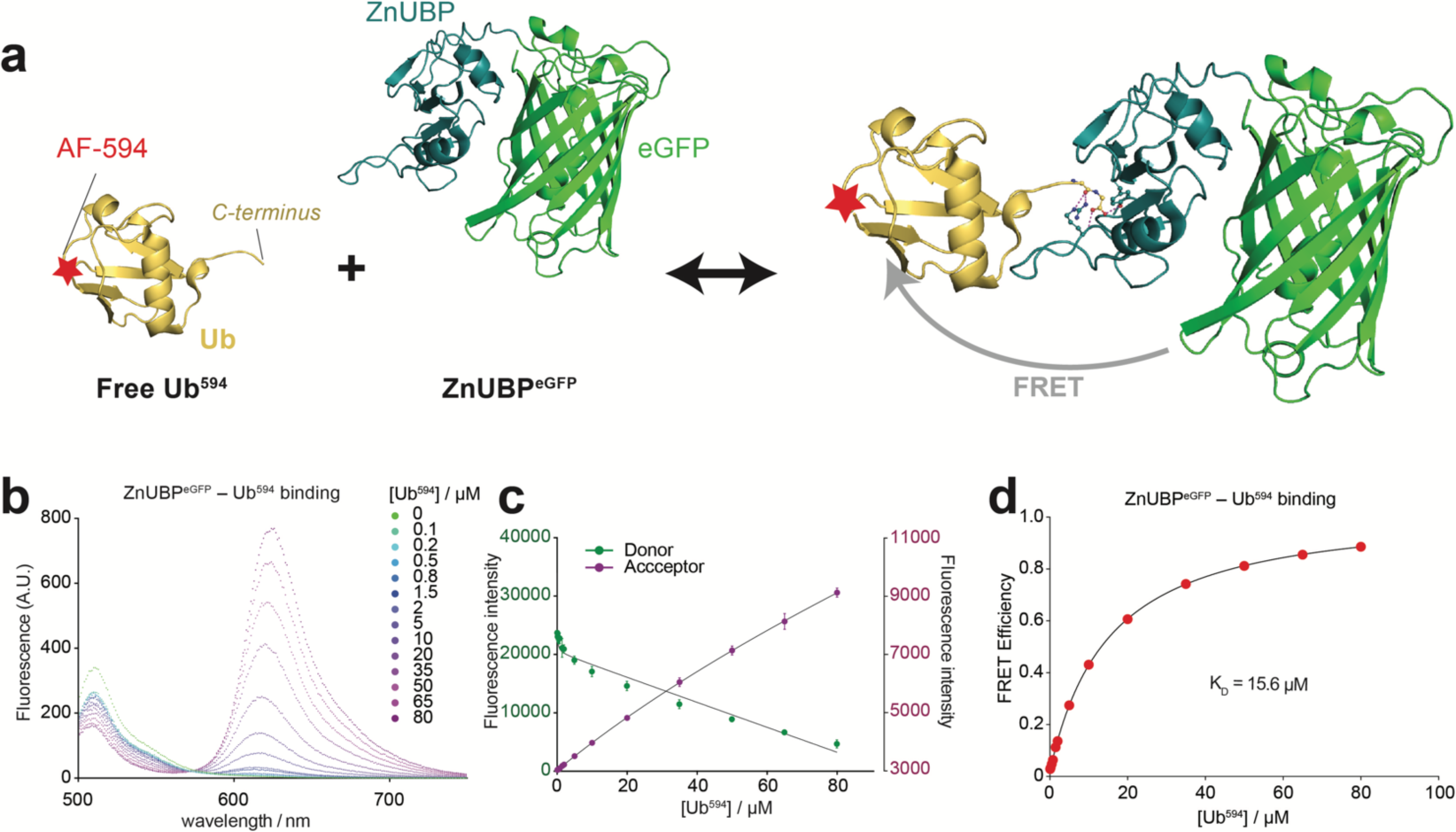
The free ubiquitin sensor system (FUSS). **(a)** A molecular model (pdb-id 2G45) of FUSS, where the Ub (yellow) is labeled at its N-terminus with an AF-594 fluorophore (red) and the ZnUBP domain (forest) is fused at its C-terminus with an eGFP (green). Specific binding between C-terminus (ball-and-stick) of the Ub to the ZnUBP is shown (*right*), enabling FRET from eGFP to AF-594. **(b)** Emission spectra (500-750 nm) collected using fluorescence excitation at 488 nm. Ub^594^ was titrated at the color-coded final concentrations into a quartz cuvette containing ZnUBP^eGFP^, continuously kept at 0.1 µM final concentration. A representative set of data from three repeats (n = 3) is shown. **(c)** Donor (green) and acceptor (scarlet) fluorescence calculated from integrated areas in **b** between 500-560 nm and 610-670 nm, respectively. **(d)** Mean FRET efficiencies calculated from the donor and emission fluorescence intensities performed in triplicate (n = 3). Error bars from standard deviations in all cases were smaller than the size of the data points (<1% of the sample mean). The binding constant is extrapolated from fitting the data to the equation for single-site binding (*equation 3*, **Materials and Methods**).

Initial rates of reaction are calculated from linear fits to the first 10 min (or the first 10 data points) after the measured start of the reaction using the equation below:

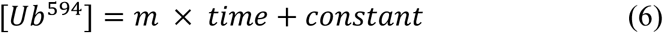

where *m* is the gradient representing the initial rate.

The Michaelis-Menten equation is defined as:

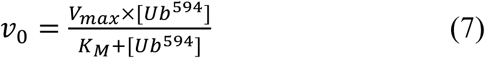

where *v*_*0*_ is the initial rate of reaction.

The Hill equation is defined as:

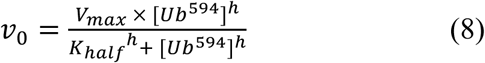

where *K*_*half*_ is the enzyme concentration at half-maximal value of *V*_*max*_ and *h* is the Hill coefficient.

### Ubiquitination system ODE simulator (USOS)

The *in silico* modeling of the ubiquitination system, USOS, is designed as follows:

1. The chain reaction is broken down into the elementary reactions (described below) that are assumed to be single-step transformations whose kinetics can be defined stoichiometrically. The chemical kinetic system is hence mathematically described using ordinary differential equations (ODE) with a set of kinetic rate laws and mass balance.
2. The species concentration versus time relationship is simulated using the ODEs, from which the initial reaction rate is calculated.
3. USOS computed rates are compared with experimental data at varying enzyme concentrations. The discrepancy between them is quantified as a loss function.
4. The model is optimized using computationally intensive machine learning method, namely simulated annealing, aided by large computing clusters. The algorithm searches for the global minimum of the loss function by running USOS over varying rate constants. The process resembles annealing where it starts at a high temperature allowing large fluctuations and slowly cools down over time until the loss function value stabilizes.
5. The optimized parameter set is plotted to identify whether the loss function converges. A *post hoc* modification of the loss function and sometimes the ODEs is conducted to direct the simulation to further convergence. The optimized parameter set then becomes the initial point for the next epoch and the above procedure is repeated until reaching an agreement between USOS prediction and data.

The set of rate laws incorporated in USOS are specified as follows:

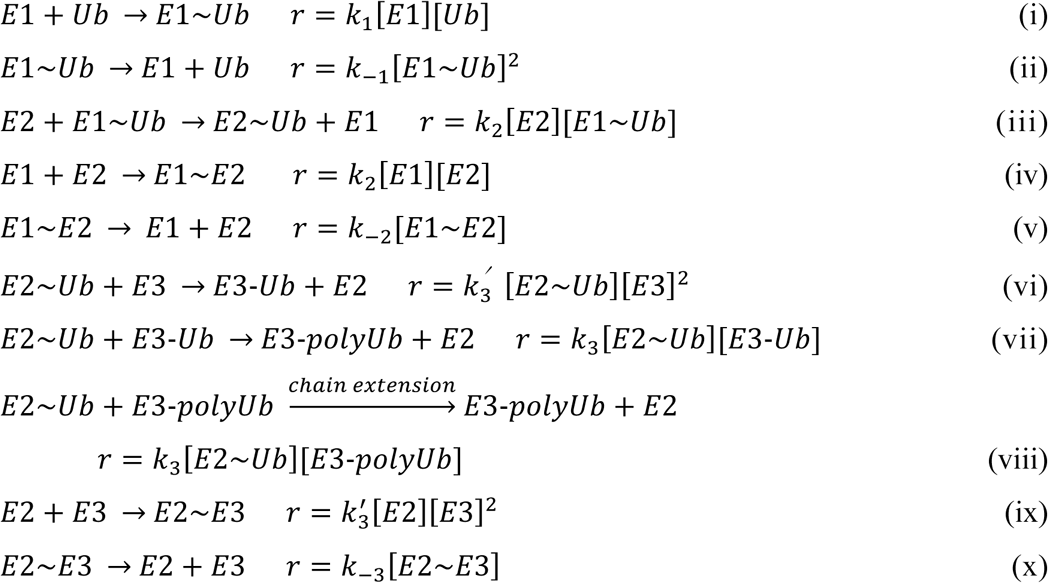

where *r* and *k* denote the reaction rate and rate constants, respectively, at the given reaction scheme. In chain extension reaction (viii) another Ub molecule is transferred from E2 to the existing Ub chain on E3, increment the polymerization degree by 1.

The MATLAB source code for USOS is available with this manuscript. It is also publicly available and can be downloaded from GitHub: https://github.com/Eric-Kobayashi/USOS.

## Results and Discussion

### A novel Free Ub Sensor System (FUSS) to study enzymatic activity

We separately expressed and purified a construct of ZnUBP domain of USP5/IsoT fused to an eGFP (ZnUBP^eGFP^) and an N-terminal Cys-modified Ub (Ub^-1Cys^) construct to apparent homogeneity. Ub^-1Cys^ was subsequently labeled with AlexaFluor(AF)-594 (Ub^594^), which is compatible with eGFP for FRET. We hypothesized that ZnUBP^eGFP^– Ub^594^ binding would establish FRET, and its efficiency would depend on the concentration of Ub^594^ in solution (**Figure 1a**). Specific recognition of Ub by ZnUBP is well-characterized and the molecular structure of the protein complex shows that the free C-terminus of Ub, which is buried inside the ZnUBP, is crucial for complex formation^31^. When conjugated, the C-terminus of Ub forms a covalent bond with other proteins and therefore would not be able to bind ZnUBP^eGFP^.

To validate our hypothesis, we titrated Ub^594^ into a quartz cuvette containing ZnUBP^eGFP^ held at a fixed final concentration (0.1 µM, see **Materials and Methods**) and measured the emission spectra from donor excitation (ex_D_ = 488 nm) in a fluorescence spectrophotometer (**Figure 1b**). Donor eGFP emission (em_D_) decreased while acceptor emission (em_A_) increased with Ub^594^ concentration (**Figure 1c**). The relationship between FRET efficiency and free Ub^594^ concentration could then be derived, with a dissociation constant (*K*_*D*_) of 15.6 µM for the binding to ZnUBP^eGFP^ (**Figure 1d**). This *K*_*D*_ is somewhat higher than previously reported (*K*_*D*_ = 2.8 µM, ^31^) and may be due to the presence of the bulky eGFP or instrumental differences. Our measured *K*_*D*_ was confirmed when repeated in a 96-well plate mode and detected by a fluorescence monochromator plate-reader (*K*_*D*_ = 15.7 µM, **Figure S1**). The optimized assay system, which we will refer to as Free Ub Sensor System (FUSS), consists of 0.1 µM ZnUBP^eGFP^ with Ub^594^ at indicated concentrations.

### FUSS detects ubiquitination and deubiquitination events in real-time

We tested whether FUSS is suitable to detect real-time ubiquitination and designed a reaction containing Uba1 (the canonical E1), UbcH5 (an E2) and CHIP (a U-box E3^32^). The ubiquitination scheme (**Figure 2a**) involving these three enzymes has been characterized before^33–35^ and is initiated upon ATP-hydrolysis and Ub-charging by Uba1^36^ (step 1), which in turn charges UbcH5 with the Ub^37^ (step 2). Charged UbcH5 (UbcH5∼Ub) may bind to dimerized and catalytic active CHIP (step 3), which readily ubiquitinates itself through a mechanism that is not fully understood^38,39^. Repeated ubiquitination ultimately results in polyubiquitinated CHIP (CHIP-polyUb, step 4). This reduces the free Ub concentration, and should thus decrease the FRET efficiency detected by FUSS. Lastly, CHIP-polyUb can be deubiquitinated by DUBs (step 5), regenerating free Ub and increasing the FRET efficiency again. Although it is possible for Uba1∼Ub or UbcH5∼Ub to discharge without their subsequent steps leading up to ubiquitination, the rates of such reactions are assumed negligible^40^ and not considered here.

**Figure 2.**
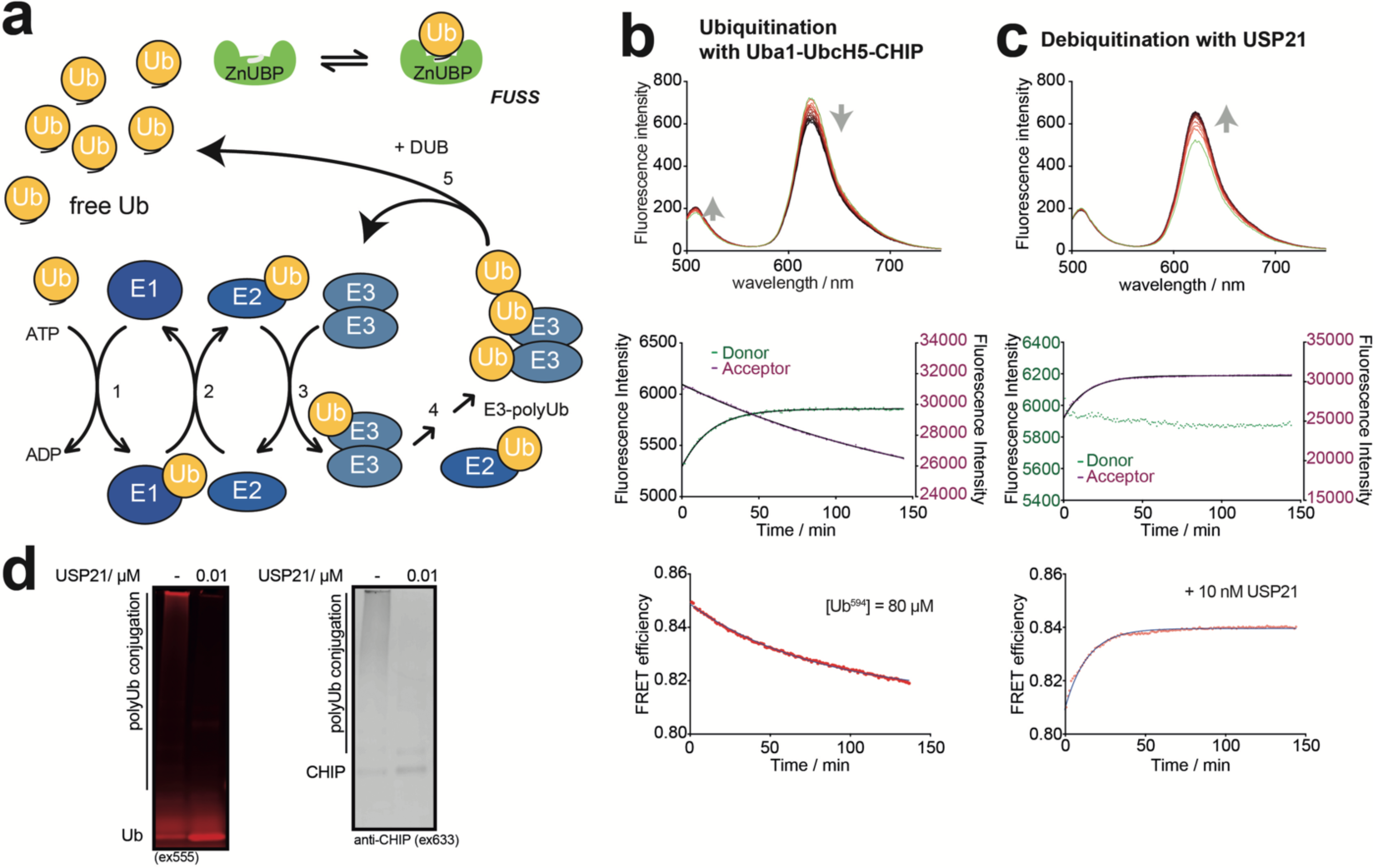
Measuring protein ubiquitination with FUSS. **(a)** A schematic representation of the major steps in Uba1-UbcH5-CHIP ubiquitination. A proportion of the free Ub^594^ pool (yellow) is bound by ZnUBP^eGFP^ (green) in a concentration-dependent manner (see **Figure 1**). In the presence of ATP, Uba1 (E1) charges free Ub^594^, forming Uba1∼Ub (*step 1*). This Ub is then transferred from Uba1 to the active site of UbcH5 (E2), leading to UbcH5∼Ub (*step 2*). UbcH5∼Ub may transfer the charged Ub to CHIP (E3, *step 3*), which ubiquitinates itself and ultimately forming CHIP-polyUb (*step 4*). Dimerization of CHIP is required for this activity, though the details of this reaction are not fully understood. Ubiquitination on CHIP can be removed by USP21 (DUB), regenerating free Ub and unmodified CHIP (*step 5*). **(b)** A ubiquitination reaction was set up in a quartz cuvette containing Uba1, UbcH5 and CHIP (0.5 µM final concentration each) mixed with FUSS ([Ub] = 80 µM). Emission spectra before (green) and after adding ATP (going from light to dark red with time) were collected continuously (*top*). Arrows indicate the direction of peak change over time. For clarity, spectra with 5 min intervals are presented. Change in donor (green) and acceptor (scarlet) fluorescence intensity over time were then calculated from the spectra (*middle*). Calculated FRET efficiency changes over time fitted with a single-exponential decay function (*bottom*). **(c)** The reaction from **b** was treated with USP21 (10 nM final concentration). Emission spectra before (green) and after (red) adding USP21 (*top*). Donor and acceptor spectra (*middle*) and the calculated FRET efficiencies (*bottom*) over time are shown as in **b**. A single-exponential decay function was used for data fitting. All experiments presented have been repeated three times (n = 3). **(d)** Completed reactions from **b** and **c** were quenched with LDS buffer containing reducing agents, separated by SDS-PAGE and detected by fluorescence scanning against Ub^594^ (*left*) or Western blot staining against CHIP using secondary antibodies labeled with AF-647 (*right*).

FUSS (Ub^594^ = 80 µM) was mixed with Uba1, UbcH5 and CHIP (0.5 µM each) and the reaction was initiated by ATP addition (**Figure 2b**, *top*). The em_D_ increased while em_A_ decreased over time (**Figure 2b**, *middle*), resulting in an overall decrease in FRET efficiency that obeyed a single-exponential decay function (**Figure 2b**, *bottom*). We then used the binding curve established in **Figure 1d** to convert FRET efficiency to Ub concentration (see *equation 4*, **Materials and Methods**), and determined the initial rate of this reaction (∼2.4 nM s^-1^).

To demonstrate the versatility of FUSS to also detect deubiquitination, USP21 was added to the cuvette after 2.5 hrs, when the ubiquitination reaction is still active. USP21 is a promiscuous DUB that is well-characterized for its high activity of deubiquitinating conjugated Ub moieties in an indiscriminate manner^29,41^, and frequently used as a generic DUB to effectively remove Ub modifications. Indeed, FRET efficiency gained over time (initial rate ∼8.8 nM s^-1^), indicating that FUSS is a reversible system (**Figure 2c**, *bottom*). Products of ubiquitination and deubiquitination were confirmed by SDS-PAGE as well as by Western blots (**Figure 2d**).

Replacing the fluorophore on Ub with AF-647 (Ub^647^), we showed that the concept of FUSS was independent of the fluorophore pair used for FRET (**Figure S2a** and **b**). We further found that in the absence of CHIP, detectable decrease in FRET efficiency could only be measured at high UbcH5 concentrations (**Figure S2c**), resulting from significant UbcH5∼Ub formation (**Figure S2d-f**). These results together suggest that CHIP-polyUb formation is the main reaction detected by FUSS.

### Determining enzyme kinetics with FUSS

The sensitivity of FUSS was tested by gradually lowering Ub^594^ starting concentrations. With decreasing Ub^594^ starting concentrations, FRET efficiency versus time plots no longer followed single-exponential functions (**Figure 3a**), and a lag-phase became apparent (∼3 min in **Figure 3b**) prior to the major decrease in FRET efficiency (reaction initiation). Such lag-phase has been observed in previous studies but not described as part of the ubiquitination and thus not taken into account in rate calculations^42^. Possibly, since Uba1 requires both Ub and ATP to initiate the reaction cascade^37^, the lag-phase suggests that lower Ub concentrations, rather than ATP, limits reaction initiation (see **Figure S3**).

**Figure 3.**
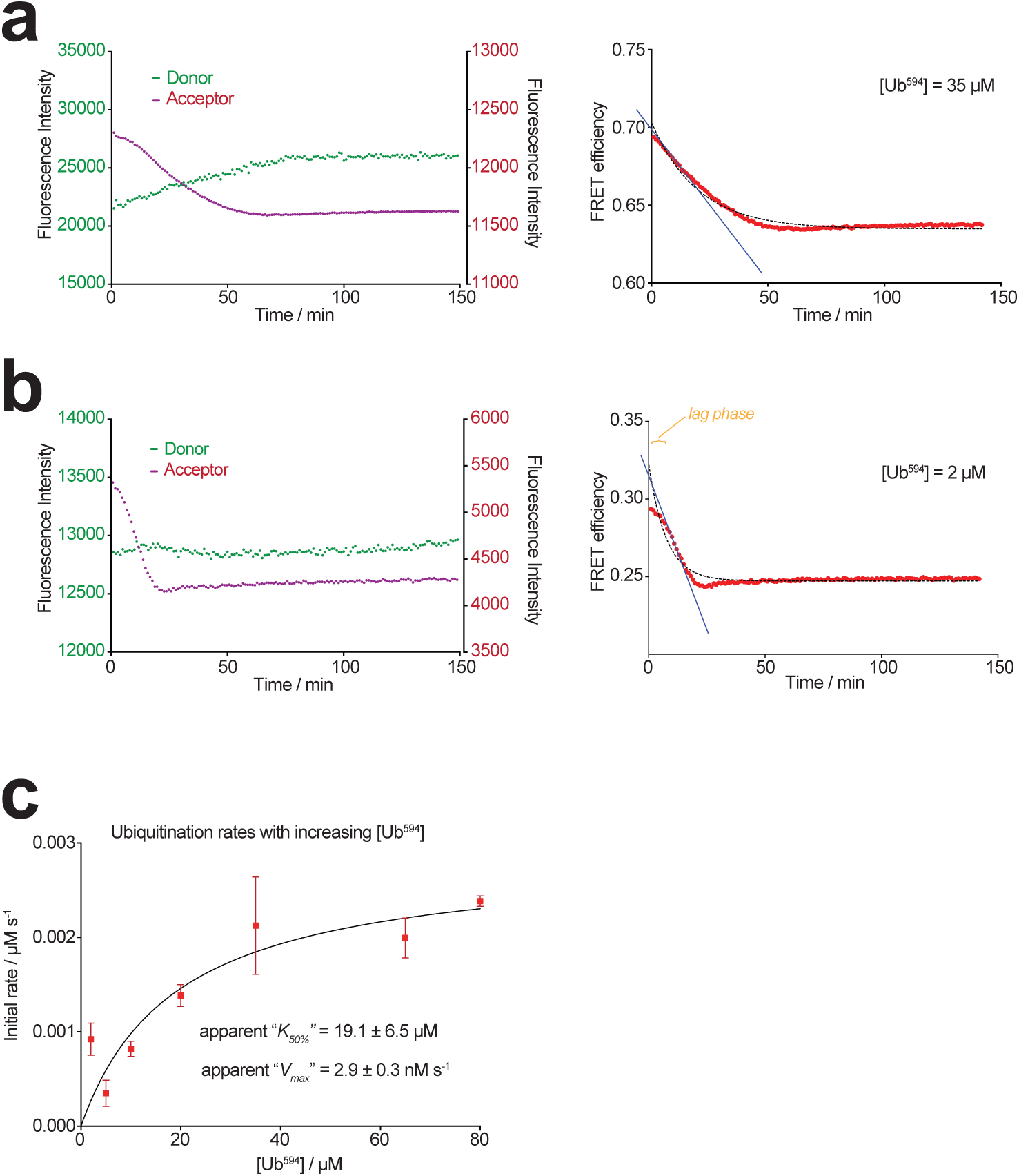
Initial rates of ubiquitination measured at various Ub^594^ concentrations. **(a)** Donor (green) and acceptor (scarlet) fluorescence intensities (*left*) and calculated FRET efficiencies (*right*) measured after ATP addition. The reaction was prepared in a quartz cuvette as in **Figure 2**, [Ub] = 35 µM. Single-exponential decay function (dotted line) does not fit the data and the initial rate was calculated from the gradient of linear fit (blue solid line) to the earlier data points. **(b)** FUSS measurement performed and presented as in **a**, [Ub] = 2 µM. The reaction ‘lag phase’ is indicated in orange (see main text). **(c)** Initial rates of reaction plotted against Ub^594^ fitted to Michaelis-Menten equation (*equation 7*) to determine apparent values of *K*_*50%*_ and *V*_*max*_. Error bars represent the standard error of the mean for three independent measurements (n = 3).

We subsequently determined the initial rates from the gradient of reactions at various Ub^594^ starting concentrations and plotted an “apparent Michaelis-Menten” curve to characterize their relationship (**Figure 3c**). The apparent *K*_*50%*_ (19.1 ± 6.5 µM) represents the concentration of Ub^594^ at which the reaction rate is half of *V*_*max*_ (2.9 ± 0.3 nM s^-1^), and conceptually equivalent to the Michaelis constant (*K*_*M*_). *K*_*M*_ is only valid when describing a scenario where Uba1, UbcH5 and CHIP would form a stable enzymatic complex, and thus not applicable here.

### FUSS is suitable for high-throughput ubiquitination assays

To demonstrate compatibility with high-throughput assays, we repeated FUSS measurements in **Figure 3** in a 96-well plate format. FRET efficiency decreased upon ATP addition compared to control (**Figure 4a**), and regained following subsequent deubiquitination with USP21 (**Figure 4b**), confirming the sensitivity of FUSS in this format. Due to the high laser exposure on the plate-reader, we further examined the em_D_ and em_A_ signals from **Figure 4a** for photo-bleaching. Significant donor photo-bleaching of eGFP^43^ was detected in the ubiquitination reaction but not in the control (**Figure 4c**). This is likely due to that most ZnUBP^eGFP^ is bound by Ub^594^ in the control, allowing the eGFP to remain photostable by transferring energy to the acceptor, AF-594. Such phenomena have been reported and experimentally validated by previous studies (e.g. ^44^). Indeed, using a different fluorophore and labeling position, we showed that the photostability of eGFP in the FUSS assay could be enhanced in a FRET-dependent manner (**Figure S4**).

**Figure 4.**
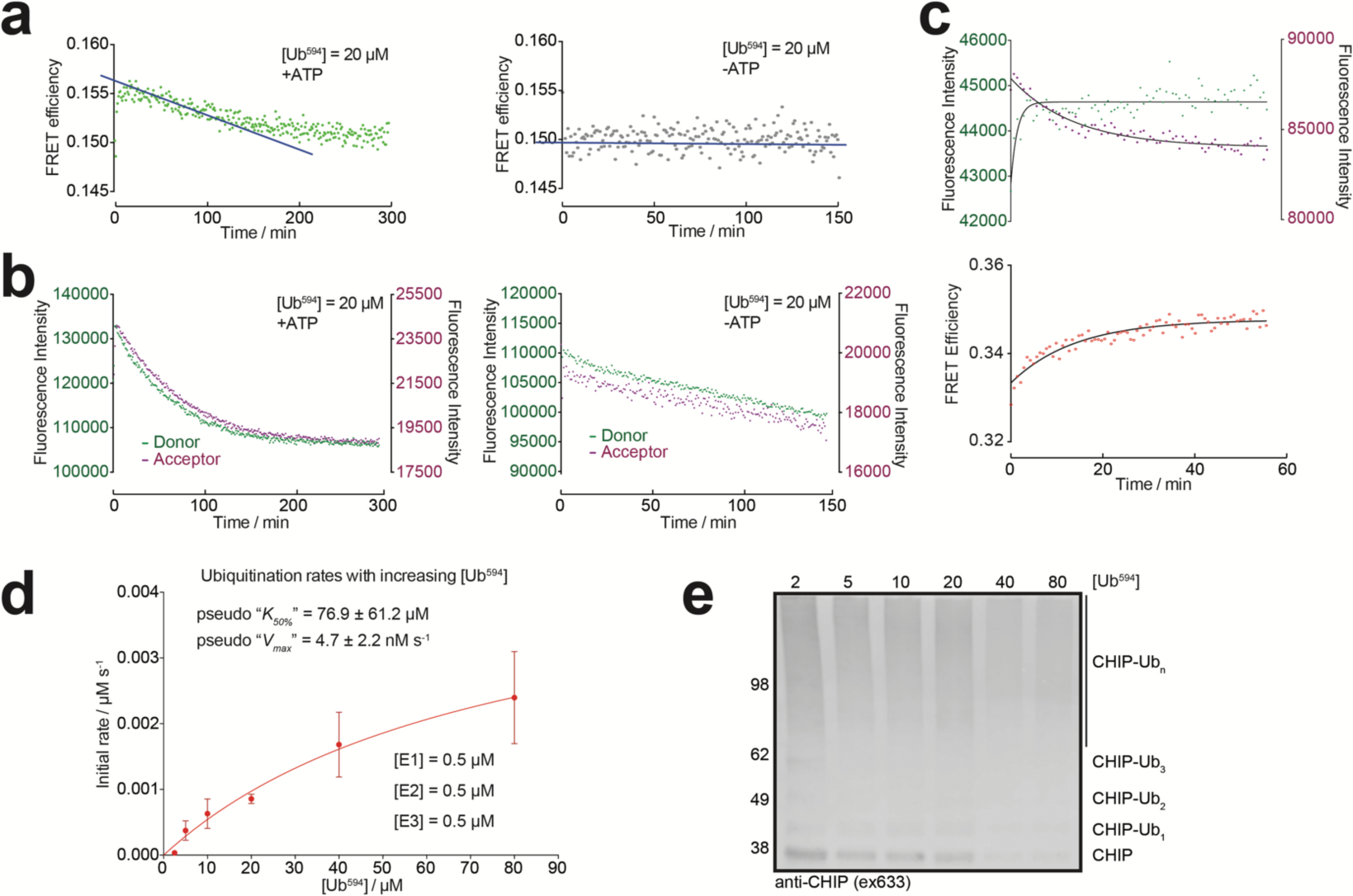
FUSS is compatible with plate-reader measurements. **(a)** Calculated FRET efficiency changes over time in a FUSS reaction ([Ub^594^] = 20 µM) containing Uba1, UbcH5 and CHIP (each at 0.5 µM) measured in a 96-well plate (*top*). The same setup is measured without adding ATP (*bottom*). **(b)** Donor (green) and acceptor (scarlet) intensities over time used for FRET calculation in **a. (c)** Detection of deubiquitinase activity in 96-well plate mode. CHIP-polyUb (ubiquitinated with 65 µM Ub^594^) was mixed with 10 nM USP21. The donor and acceptor intensities (*top*) and the calculated FRET efficiencies (*bottom*) are shown. **(d)** Initial rates of reaction extracted from **Figure S6** (calculated using *equation 5* in **Materials and Methods**) were fitted to the Michaelis-Menten equation (*equation 7*). Error bars represent the standard error of the mean for three repeat measurements (n = 3). **(e)** Anti-CHIP Western blot of the final reaction products in **d** quenched with LDS buffer containing reducing agent. The various forms of CHIP are indicated. The secondary antibody was labeled with AF-647 and the blot was detected using a Typhoon fluorescence scanner (see **Materials and Methods**). Numbers to the left represent molecular weight markers.

Given that eGFP photo-bleached rapidly upon Ub^594^ dissociation, we calculated FRET efficiencies using pre-determined em_D_ values from Ub^594^-associated ZnUBP^eGFP^. Since the same final concentration of ZnUBP^eGFP^ (0.1 µM) was used in all FUSS reactions, we repeatedly measured ZnUBP^eGFP^ bound to 5 µM Ub^594^ and calculated an average em_D_, which remained stable over time (**Figure S5**, standard deviation < 15%, n = 6). This average em_D_ was substituted into FRET efficiency calculations with the measured em_A_, and allowed comparisons between reactions performed on the plate-reader.

We subsequently used the plate-reader to determine the apparent *K*_*50%*_ (76.9 ± 61.2 µM) and *V*_*max*_ (4.7 ± 2.2 nM s^-1^) from the initial rates of reaction of Uba1-UbcH5-CHIP (each at 0.5 µM final concentration) at increasing Ub^594^ concentrations (**Figure 4d** and **Figure S6a**). As expected, the final reaction products contained a higher level of CHIP-polyUb with increasing Ub concentrations (**Figure 4e** and **Figure S6b**).

### Increasing E2 or E3 concentrations enhances initial ubiquitination rates

Having established a high-throughput approach, we next investigated how enzyme concentrations influenced ubiquitination rates. Increasing concentrations of either Uba1, UbcH5 or CHIP were mixed with the FUSS ([Ub^594^] = 5 µM) containing the other two enzymes both fixed at 0.5 µM final concentration. We first determined the initial rates of reaction at increasing CHIP concentrations, which showed a sigmoidal relationship (excluding the highest [CHIP]) in a semi-log plot that reached its maximum (0.9 ± 0.1 nM s^-1^) when [CHIP] > 2 µM (**Figure 5a**). This appeared to be concurrent with a changing pattern of the final CHIP-polyUb products from high molecular weight (MW) smears to distinct ubiquitinated CHIP bands between 40-100 kDa (**Figure 5b** and **Figure S7a**).

**Figure 5.**
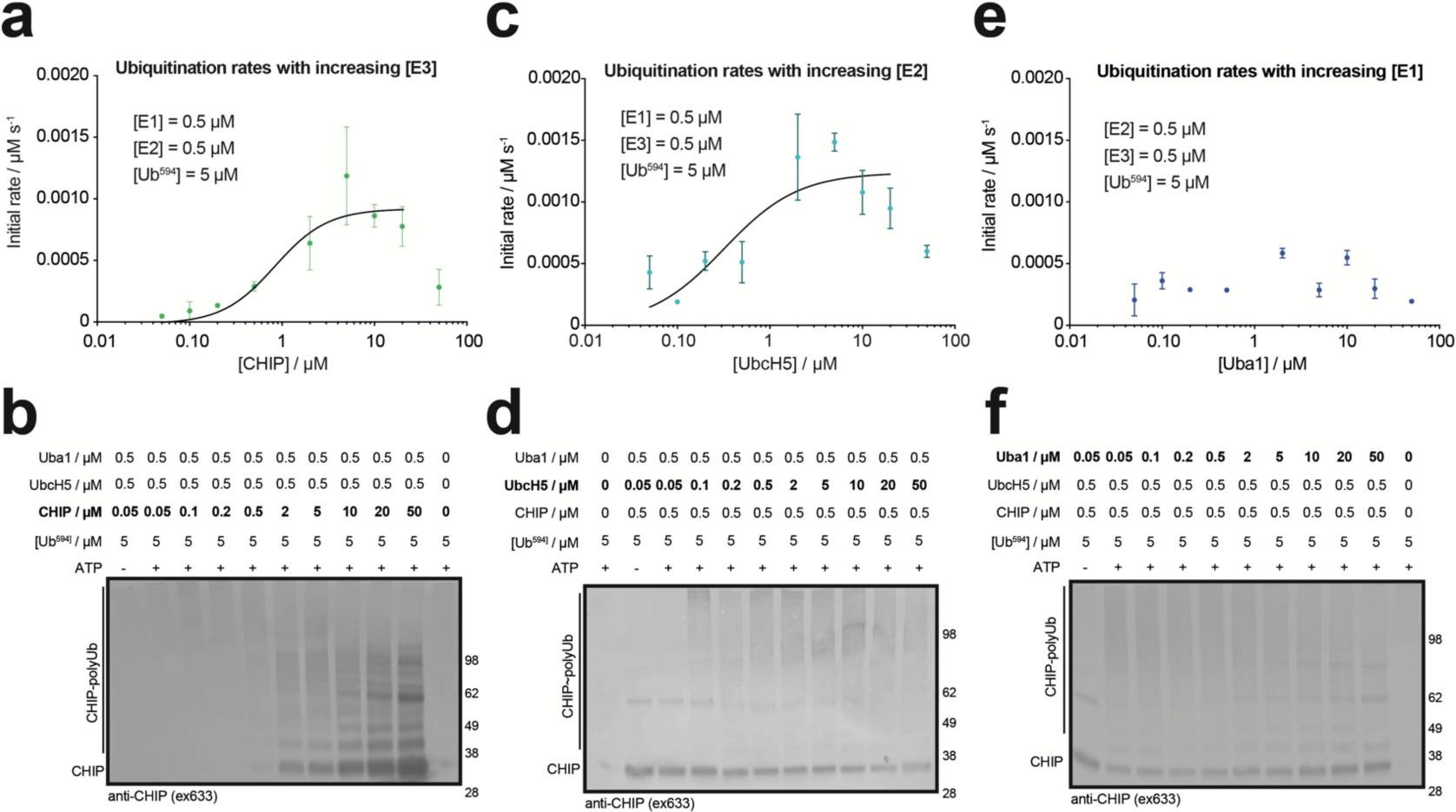
Relationship between enzyme concentration and ubiquitination rate. **(a)** CHIP at increasing concentrations was incubated with FUSS reactions ([Ub^594^] = 5 µM) containing Uba1 and UbcH5 at 5 µM each. The initial rates of reaction are shown in a semi-log plot against CHIP concentration and fitted with the Hill equation, giving Hill coefficient = 1.5 ± 0.6. **(b)** Western blot against CHIP from reactions in **a** after 150 min. A sample of uncatalyzed reaction (lane 1) and a sample of Ub^594^ only (lane 11) were included as controls. All samples were quenched with LDS buffer containing reducing agent. **(c)** UbcH5 at increasing concentrations was incubated with FUSS reactions ([Ub^594^] = 5 µM) containing Uba1 and CHIP at 5 µM each. Data points were fitted as in **a**, giving Hill coefficient = 1.1 ± 0.6. **(d)** Anti-CHIP Western blot of the reactions in **c** after 150 min, performed as in **b. (e)** Uba1 at increasing concentrations was incubated with FUSS reactions ([Ub^594^] = 5 µM) containing UbcH5 and CHIP at 5 µM each. **(f)** Anti-CHIP Western blot of the reactions in **e** after 150 min. Error bars represent the standard error of the mean for three repeat measurements (n = 3).

Similarly, changes in the initial rate with increasing UbcH5 concentrations could also be described with a sigmoidal relationship (excluding the highest [UbcH5]). The maximum rate of ubiquitination (1.1 ± 0.1 nM s^-1^) was reached at [UbcH5] = 5 µM (**Figure 5c**), consistent with **Figure 5a**. As with CHIP, very high UbcH5 concentrations did not appear to support further increases in the rate, and the high MW smear of ubiquitination products remained at very high UbcH5 concentrations (**Figure 5d** and **Figure S7b**).

In contrast, increasing Uba1 concentration had little effect on ubiquitination rates (**Figure 5e**) and the maximum rate at 0.5 ± 0.06 nM s^-1^ (at [Uba1] = 2 µM) was much lower than those in **Figure 5a** and **c**. This suggests that the initial rate of ubiquitination cannot be accelerated by increasing Uba1 concentration. As found at very high CHIP and UbcH5 concentrations, the level of ubiquitination products also decreased at very high Uba1 concentrations (**Figure 5f** and **Figure S7c**).

Possibly, reduced level of ubiquitination at high enzyme concentrations could be due to competition between charged and uncharged enzymes for the same binding sites, so that the Ub-charged enzyme cannot transfer its Ub moiety down the E1-E2-E3 pathway. For instance, at high concentrations most of UbcH5 will remain uncharged ([UbcH5] > [Ub] = 5 µM), but still able to bind CHIP^39^ and compete with UbcH5∼Ub for CHIP interactions. The rate would therefore reduce as [UbcH5] increases beyond 5 µM. An additional source of the reduced ubiquitination rate could be due to futile cycles of Ub-charging and discharging without Ub transfer to the enzyme downstream, reportedly influenced by the level of enzymes present^40^. To further validate these experimental observations, we applied *in silico* modeling with a machine learning approach for the reactions in **Figure 5** using parameters inferred from the experimental data (see **Materials and Methods**). Our modeled results are generally in agreement with our experimental observations, supporting reduced ubiquitination rates at very high enzyme concentrations (**Figure S8**).

### Rate of ubiquitination (nM s^-1^) increases in the presence of tau

Under physiological conditions CHIP ubiquitinates misfolded proteins and aggregates and targets these for degradation^33,45^. One substrate of CHIP is tau^46,47^, an amyloidogenic protein associated with Alzheimer’s disease^48^. We assessed whether the presence of this true substrate would alter ubiquitination rates. Initial rates of ubiquitination (in nM s^-1^) increased in the presence of tau (**Figure 6a**), and the final Ub-modified tau product was confirmed by Western blot (**Figure 6b**). The level of CHIP-polyUb formed appeared to remain largely the same whether in the presence or absence of tau (**Figure 6c**). Together, these results suggest that tau potentially increases the overall activity of CHIP, which readily modifies both itself and its substrate.

**Figure 6.**
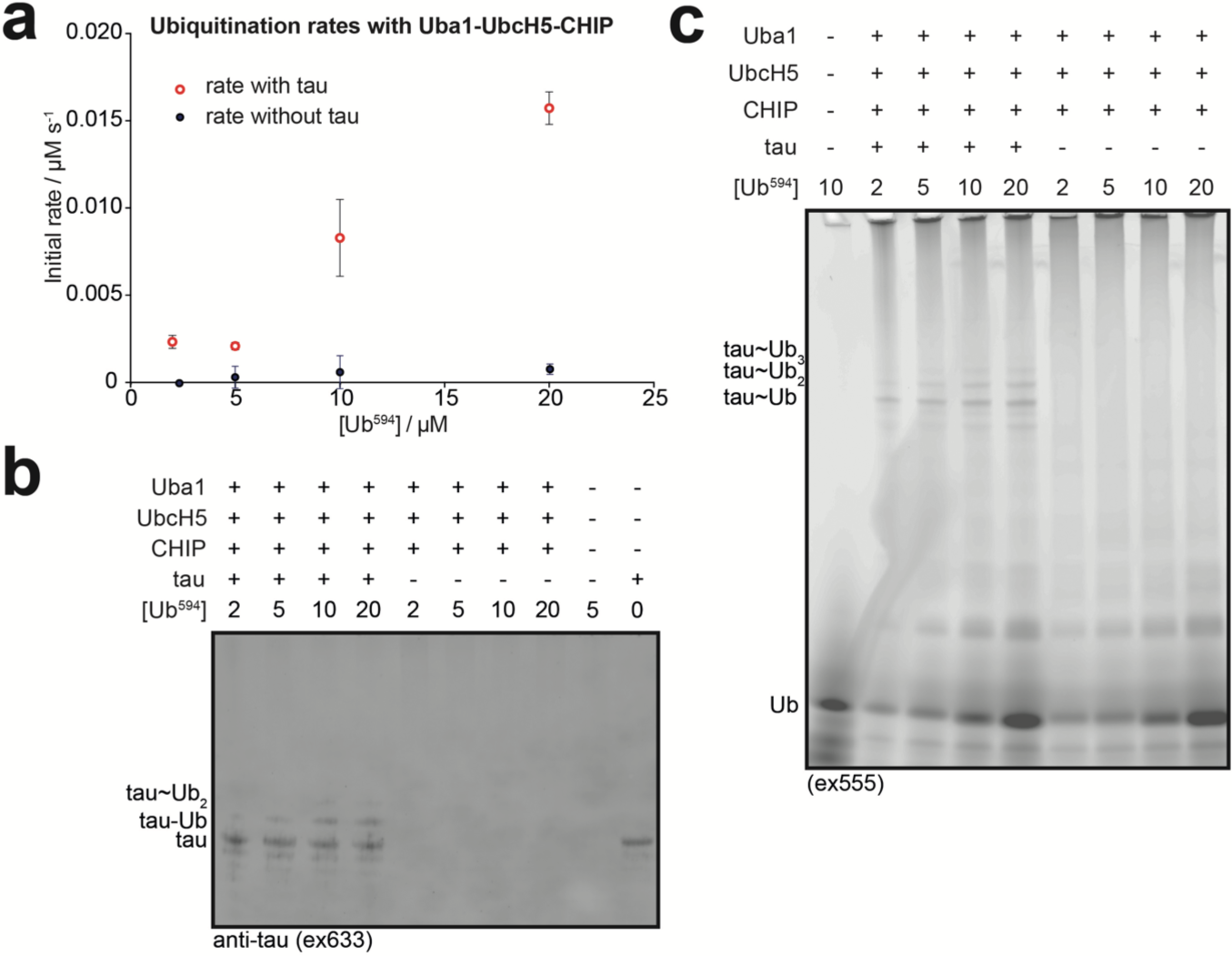
Adding tau increases the rate of ubiquitination. **(a)** Uba1, UbcH5 and CHIP (5 µM each) with or without 10 µM tau was mixed with FUSS reactions ([Ub^594^] at 2, 5, 10 or 20 µM) and the initial rates were plotted against Ub^594^ concentrations. **(b)** Anti-tau Western blot of final samples of reaction from **a** with tau (lanes 1-4) and without tau (lanes 5-8) run next to each other. A sample of tau only (lane 10) is included as a control. **(c)** Fluorescence scan against Ub^594^ of the samples in **b** separated by SDS-PAGE. A sample of Ub^594^ only (lane 1) is included as a control. All samples were quenched with LDS buffer containing reducing agent before separation on SDS-PAGE.

## Conclusions

We have demonstrated in this study a novel assay, FUSS, to determine enzyme kinetics by sensitively measuring changes in the concentration of free Ub in solution. Because FUSS is not ‘fussy’ about the enzymes involved in ubiquitination, some E2s that directly ubiquitinate substrates (e.g. ^49,50^) without assistance from E3s would also be detected by FUSS. Depending on the nature of investigation, FUSS may be further modified to e.g. introduce modifications on the Ub moiety to prevent discharging from E2 or E3s (e.g. ^51,52^), thus enabling characterization of distinct ubiquitination steps (see **Figure 2a**). A second feasible application may be to determine binding affinity for UbIPs using FUSS, which would show reduced FRET as the UbIP is titrated into the solution^22^. Finally, wild-type Ub (Ub^WT^) could be mixed and used to compete with Ub^594^ for binding with ZnUBP^eGFP^, reducing the FRET signal. In line with this idea, FUSS may be further engineered so that the binding of Ub^594^ would be weaker than Ub^WT^ and at the same time unable to be charged by E1. Any ubiquitination will then result in binding of Ub^594^ to ZnUBP^eGFP^ that were previously occupied by Ub^WT^, increasing the FRET signal. This approach could be especially useful in measuring ubiquitination or deubiquitination activities in e.g. cell lysates. Together, our FUSS assay offers a simple proof-of-principle approach to study the enzymes of the Ub system and enable the wider scientific community interested in the Ub system to study its enzymes.

## Acknowledgement

YY conceptualized research, prepared reagents, performed experiments, analyzed the data and wrote the manuscript. BKC measured enzymatic reactions under supervision of YY. YZ developed the USOS model and applied machine learning to the *in silico* simulation design. KJ purified proteins and performed experiments. DF and DK provided reagents. YY is supported by a Fellowship from UK Dementia Research Institute, which receives its funding from UK DRI Ltd, funded by the UK Medical Research Council, Alzheimer’s Society and Alzheimer’s Research UK. Work in DF lab is funded by NIH grant R01-GM043601. This project was funded by a Sir Henry Wellcome Fellowship awarded to YY.

## Supporting information

The following files are available free of charge.

Supporting information (Zuo_etal_SI.pdf)

USOS source code (USOS.zip)

