## Supplementary Information for "A general *in vitro* assay to study enzymatic activities of the ubiquitin system"

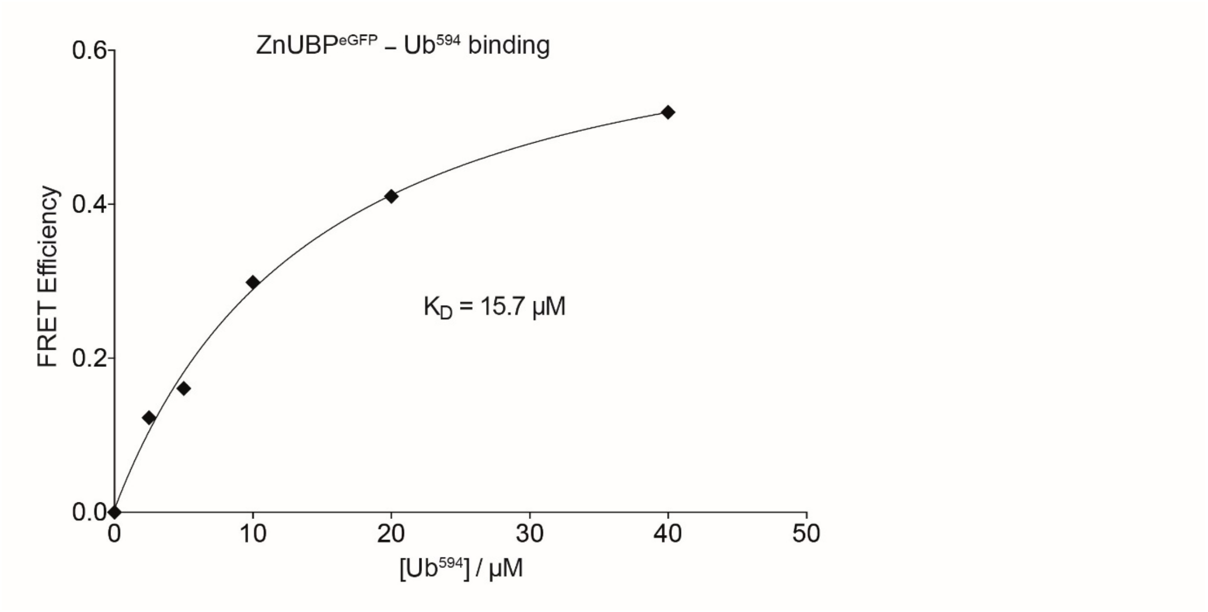


**Figure S1.** The relationship between FRET efficiency and Ub concentration measured on a plate-reader. Mean FRET efficiencies were calculated from the measured donor and emission fluorescence intensities performed in triplicate (n = 3). Error bars representing standard deviations in all cases were smaller than the size of data points (<1% of mean). The dissociation constant ($K_D$) is extrapolated from fitting the data to the equation for single-site binding (*equation 3*, **Material and Methods**).

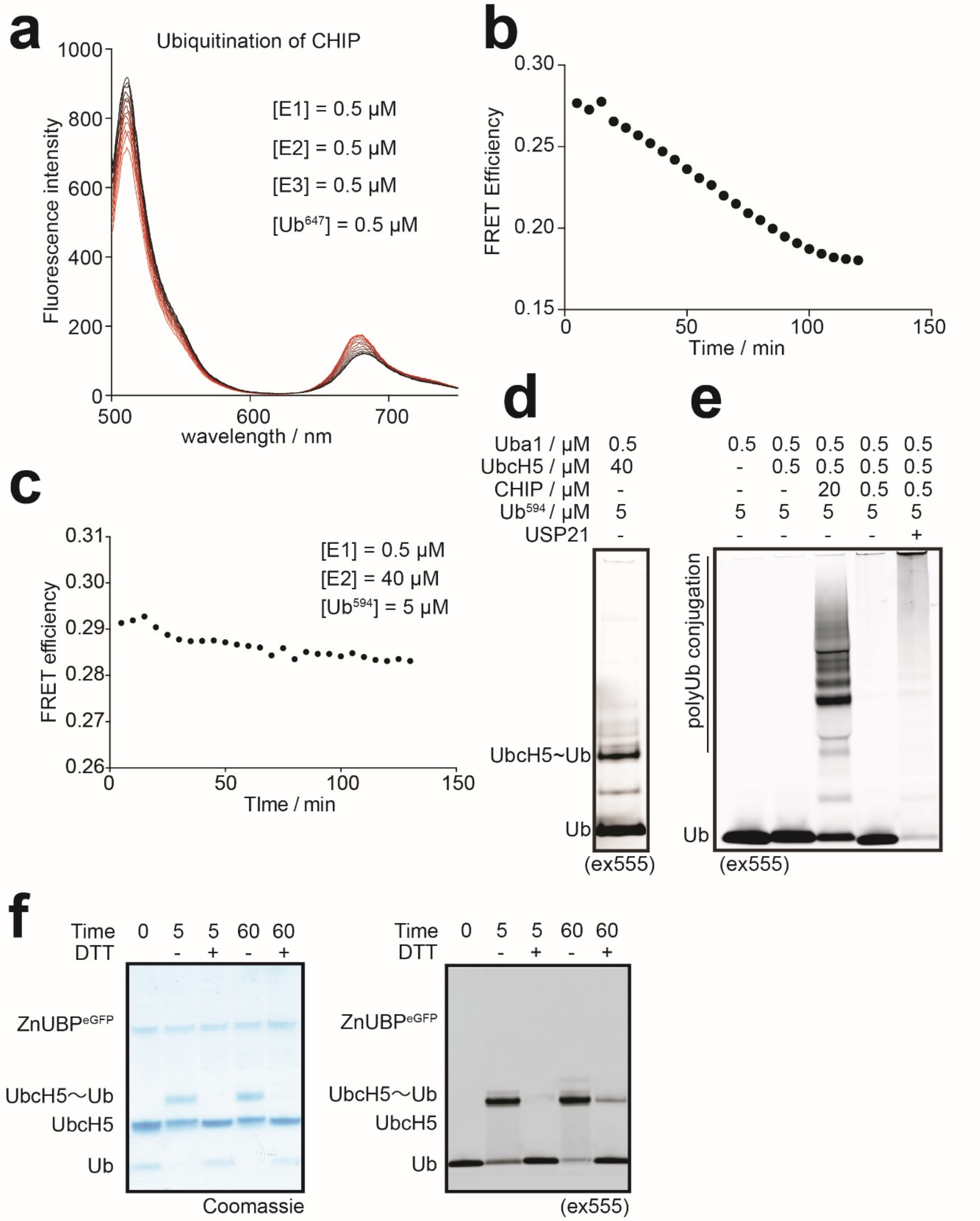


**Figure S2.** Polyubiquitination of CHIP. **(a)** FUSS reaction was set up in a cuvette with $Ub^{647}$ at 0.5 µM and Uba1, UbcH5 and CHIP (0.5 µM final concentration each). Measurements were performed as in **Figure 2**, with 5 min interval time between measurements. **(b)** Calculated FRET efficiencies from the data in **a**. **(c)** A ubiquitination reaction containing Uba1 at 0.5 µM and $Ub^{647}$ at 5 µM required high UbcH5 concentration (40 µM) for FUSS to detect a change in FRET efficiency. **(d)** Product of the reaction in **c** separated by SDS-PAGE and scanned on typhoon detects UbcH5~Ub conjugates. **(e)** Ubiquitination reactions containing FUSS ($Ub^{647}$ at 5 μM) mixed with Uba1 only (lane 1), Uba1 with UbcH5 (lane 2) or Uba1, UbcH5 and CHIP at high- (lane 3) or low concentrations (lane 5). The ubiquitinated products in lane 5 are deubiquitinated by USP21 (lane 4). All reactions were incubated for 2 hrs and quenched with LDS buffer containing reducing agent before separation by SDS-PAGE, followed by visualization on a Typhoon scanner. **(f)** Control samples of charged UbcH5~Ub formed over time, which appear to be deconjugated by DTT on SDS-PAGE and Coomassie detection, but fluorescence scan reveals residual UbcH5~Ub in the 60 min reaction sample.

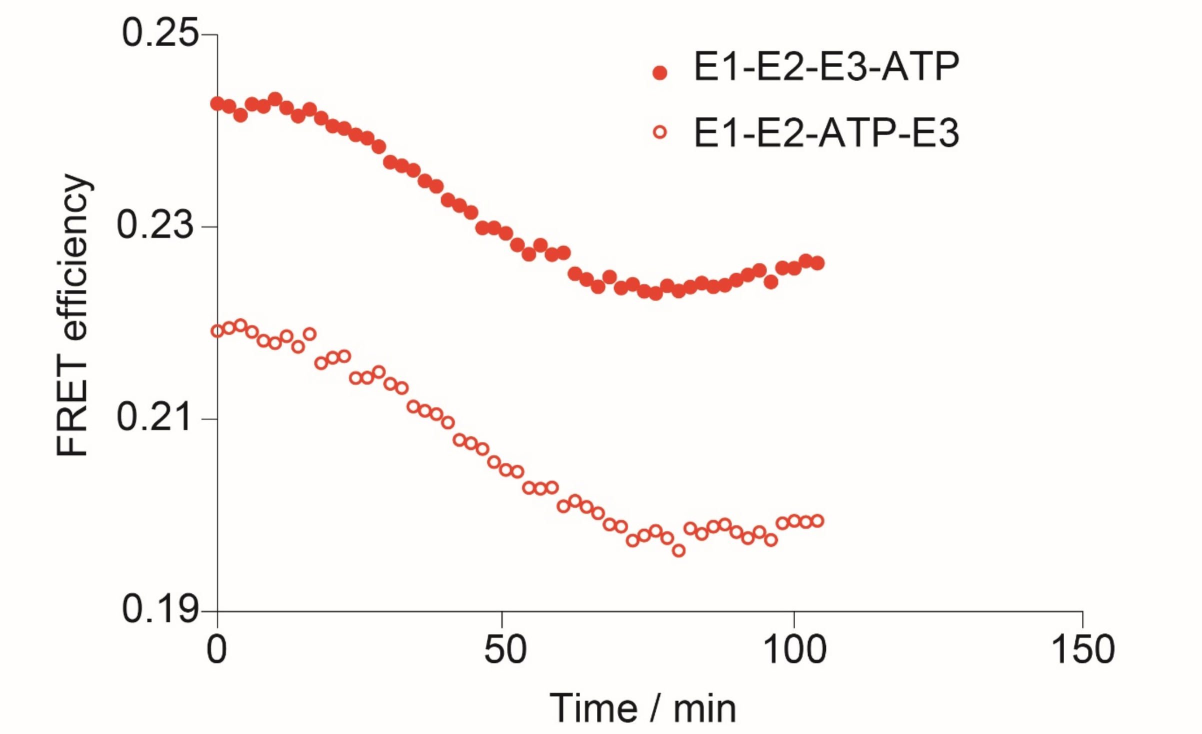


**Figure S3.** The rate of reaction is not limited by ATP hydrolysis. Two reactions with identical final composition were simultaneously measured in quartz cuvettes after all components were added (as described in **Figure 3**). One reaction was pre-incubated with ATP (10 mM final concentration) for 5 min before adding CHIP ('E1-E2-ATP-E3'), to allow pre-charging of Uba1~Ub and UbcH5~Ub before CHIP ubiquitination occurs. The second reaction was pre-incubated with all three enzymes (0.5 μM each) for 5 min before adding ATP ('E1-E2-E3-ATP'), the typical procedure used in the current study. The experiment was repeated independently three times (n = 3), with a typical set of ubiquitination reaction shown here.

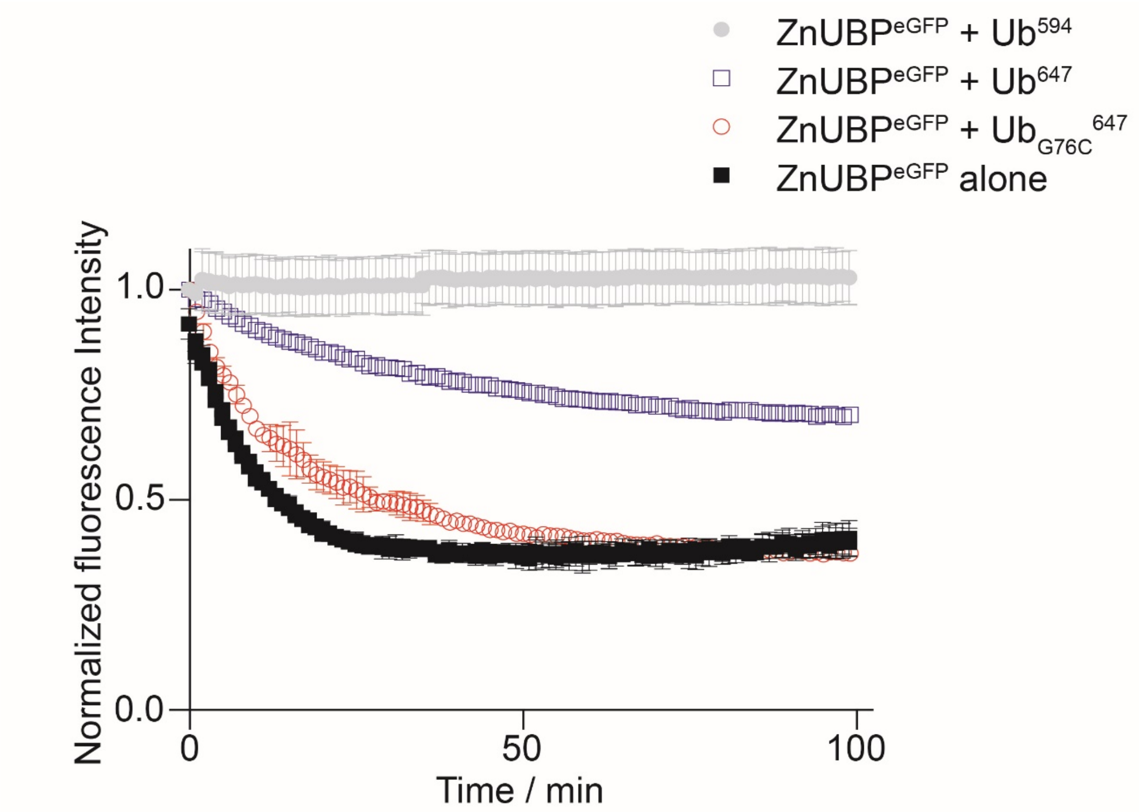


**Figure S4.** Donor photo-bleaching is dependent on FRET efficiency. Fluorescence intensity from donor ZnUBP$^{eGFP}$ emission is stable when bound to Ub$^{594}$ (grey). Replacing the fluorophore with AF-647 (blue) decreases the efficiency of energy transfer from ZnUBP$^{eGFP}$, resulting in a somewhat faster decay of donor fluorescence. Coupling the AF-647 fluorophore to the C-terminus of Ub mutant Gly76Cys (Ub$_{G76C}$) prevents effective binding to ZnUBP$^{eGFP}$ (red), resulting in rapid decay of the donor fluorescence, comparable to ZnUBP$^{eGFP}$ alone (black).

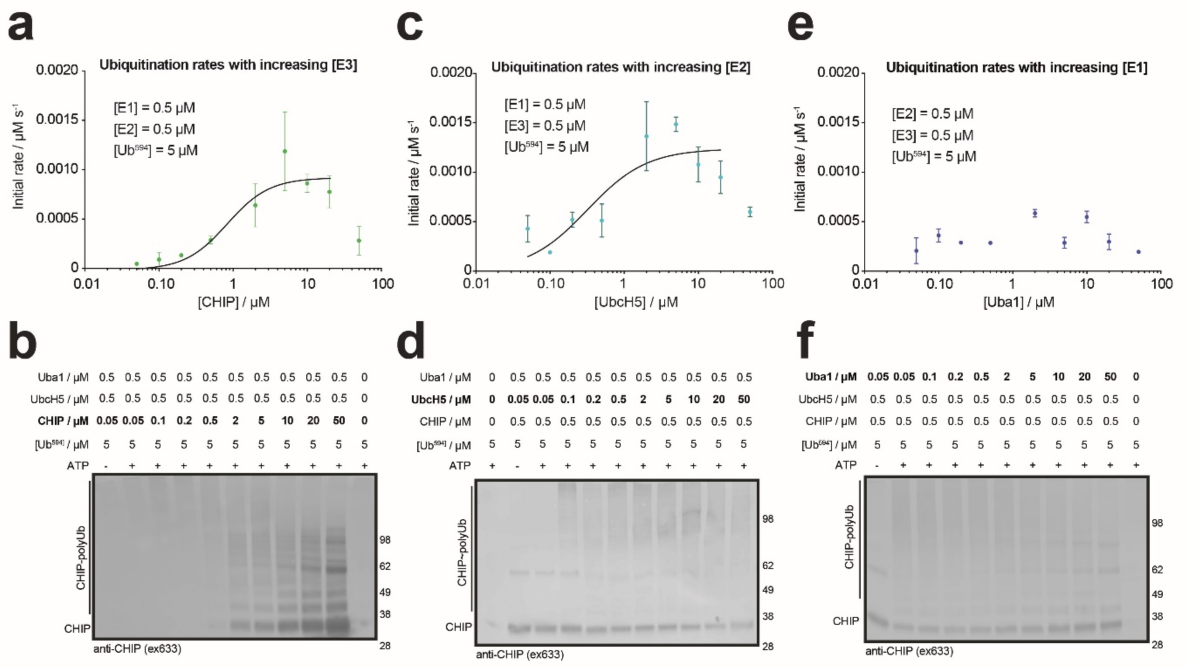


**Figure S5.** Donor fluorescence does not suffer from photo-bleaching when bound to Ub$^{594}$. Mean values of six independent measurements of ZnUBP$^{eGFP}$ (0.1 µM) bound to 5 µM Ub$^{594}$ and measured over time on the plate-reader. Error bars represent standard deviation (n = 6). **(b)** FUSS assay ([Ub$^{594}$] = 5 µM) set up as in **Figure 3** to acquire donor and acceptor emissions. Donor fluorescence does not change at this Ub$^{594}$ concentration even as ubiquitination reaction continues. **(c)** Calculated FRET efficiency from donor and acceptor intensities in **b**.

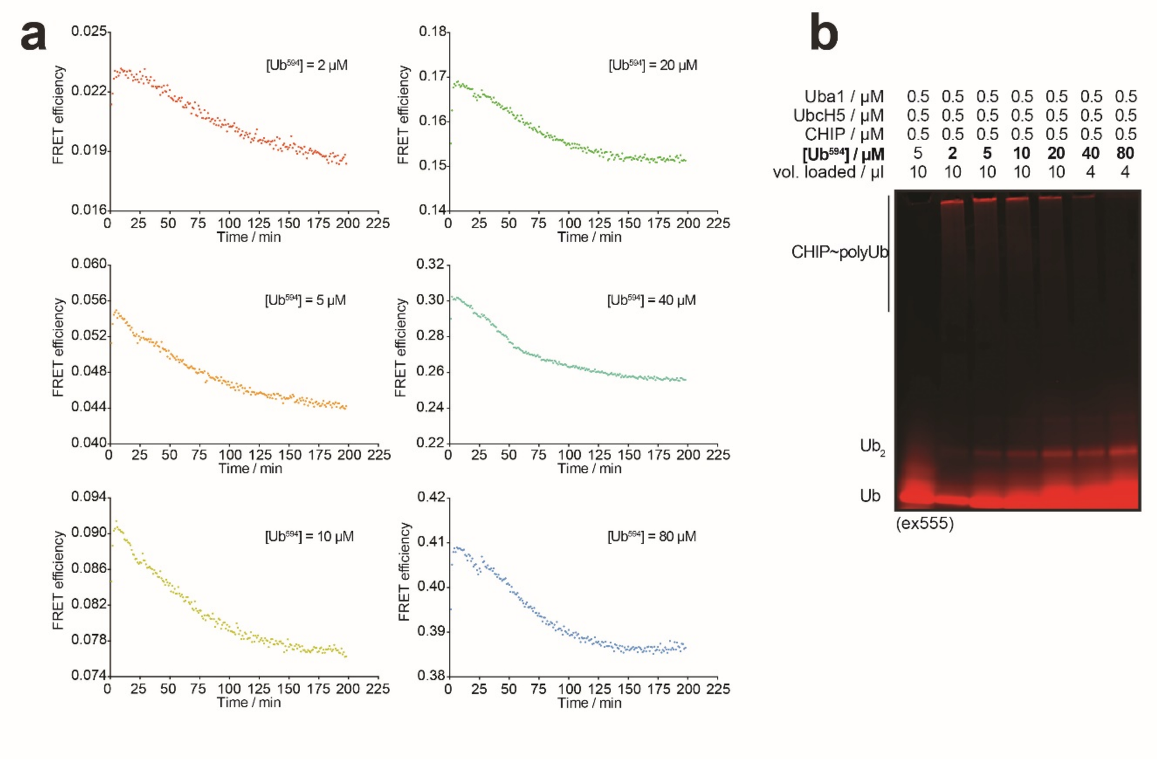


**Figure S6.** Plate-reader measurement of ubiquitination reactions at increasing $Ub^{594}$ concentrations. **(a)** FRET efficiencies were calculated from measured emission intensity and the donor fluorescence in **Figure S5a**. $Ub^{594}$ concentrations (2, 5, 10, 20, 40 and 80 μM) used in FUSS assays are indicated on each plot. Representative data of three repeat measurements (n = 3) are shown here. **(b)** Samples from completed reactions in **a** were quenched with LDS buffer containing reducing reagent, separated by SDS-PAGE and scanned on a typhoon scanner to detect ubiquitinated bands.

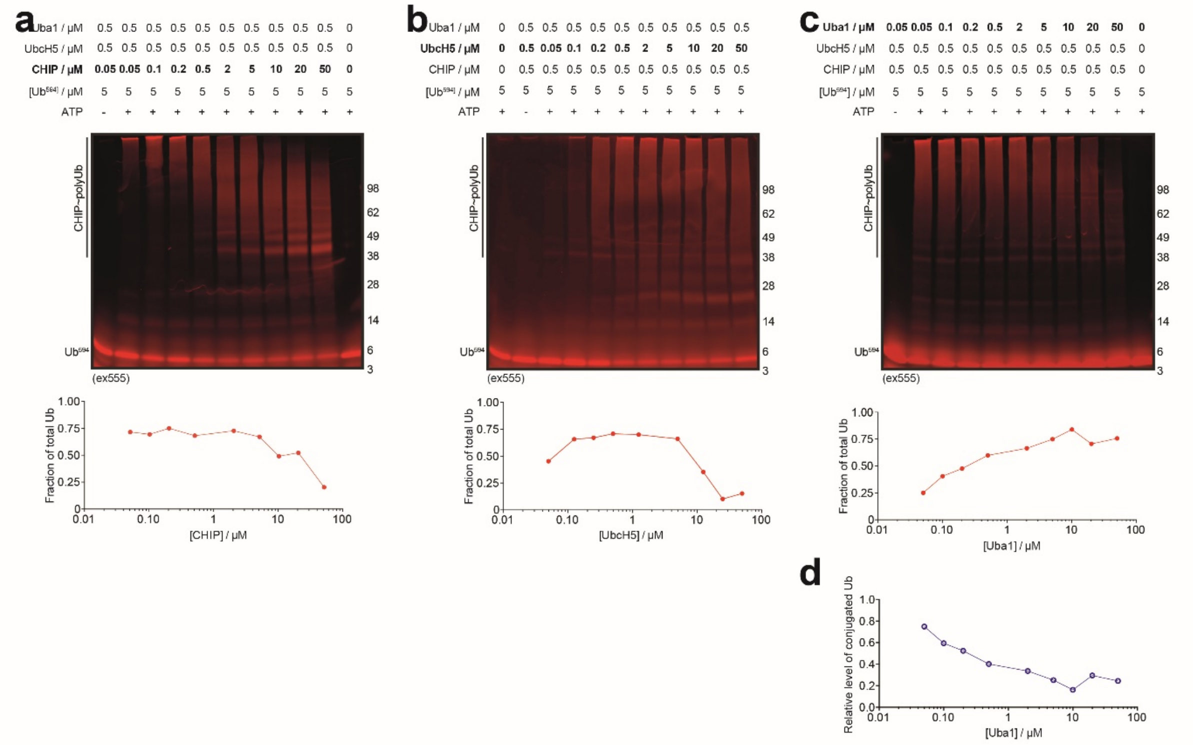


**Figure S7.** Ubiquitination pattern changes with enzyme concentration. Completed reactions from **(a)** CHIP concentration gradient in **Figure 5b**, **(b)** UbcH5 concentration gradient in **Figure 5d,** and **(c)** Uba1 concentration gradient in **Figure 5f** were separated by SDS-PAGE and scanned on Typhoon to detect ubiquitinated bands. The final concentrations of each protein are indicated on the top of every lane. All samples were quenched with LDS buffer containing reducing agent. The fraction of free Ub in each lane is quantified from the ratio of intensity compared to control lane (Ub only) and shown on the bottom of each gel. **(d)** The trend between fraction of conjugated Ub and Uba1 concentration suggests a negative correlation: the higher the enzyme concentration, the smaller the fraction of Ub that are conjugated to other proteins.

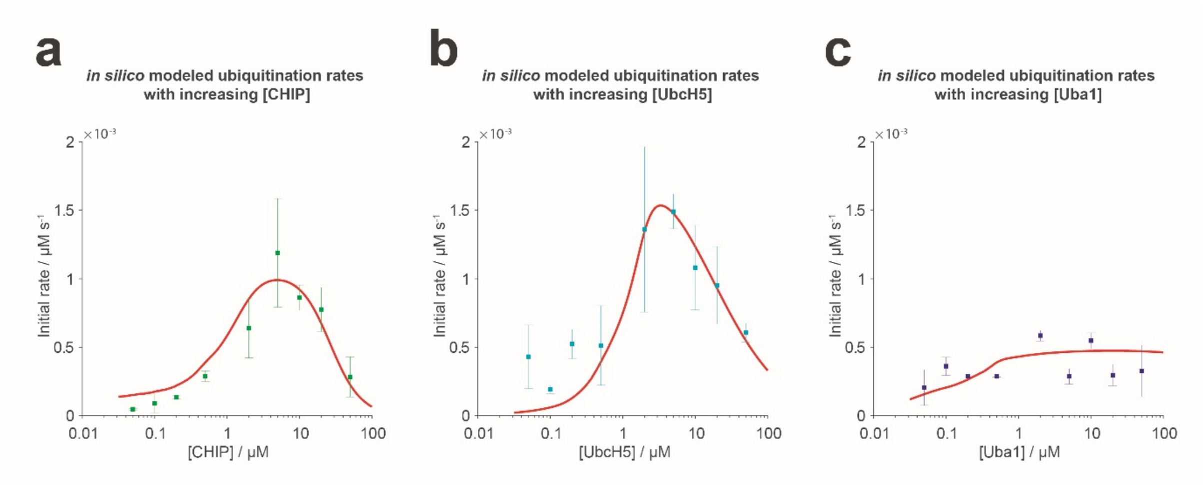


**Figure S8.** *in silico* simulation of the ubiquitination chain reaction. This model, named ubiquitination system ODE simulator (USOS), was generated by combining a rational design of ODEs (ordinary differential equations) as the kinetics framework and the computational inference of the ODE parameters, i.e. the rate constants, from the experimental data using a machine learning approach (see **Materials and Methods**). A close fit can be observed between USOS simulated results (red solid line) and the experimental data from **Figure 5** (square points) at varying enzyme concentrations of **(a)** CHIP; **(b)** UbcH5 and **(c)** Uba1. This close fit suggests that the mechanism described by USOS is highly probable. The machine learning approach gives the following rate constants (as defined in Materials and Methods for reaction schemes i-x): $k_1 = 987.6$, $k_2 = 1.880 \times 10^7$, $k_3 = 1.235 \times 10^3$, $k_3' = 2.470 \times 10^9$, $k_{-1} = 6.105 \times 10^{10}$, $k_{-2} = 320.6$, $k_{-3} = 0.8933$. All units are given in $M^{-1} s^{-1}$, apart from $k_3'$, which is in $M^{-2} s^{-1}$. The dissociation reaction of E1~Ub ($k_{-1}$) has a very high rate, potentially suggesting that the activation by E1 could be the overall rate-limiting step. A largely unchanged reaction rate with increasing [E1] can be explained by the rapid dissociation of E1~Ub, rather than the slow association between E1 and Ub, given the modeling results. It further suggests that the E2~E3 enzyme complex is stable ($k_{-3}$ is small), thus the reaction rate goes down at very high [E2] or [E3]. USOS also recognizes that E3 exists in the solution as a dimer, therefore the reactions involving E3 (vi and ix, see **Materials and Methods**) are quadratically dependent on [E3].
